# Genetic and phylogenetic uncoupling of structure and function in human transmodal cortex

**DOI:** 10.1101/2021.06.08.447522

**Authors:** Sofie L. Valk, Ting Xu, Casey Paquola, Bo-yong Park, Richard A.I. Bethlehem, Reinder Vos de Wael, Jessica Royer, Shahrzad Kharabian Masouleh, Şeyma Bayrak, Peter Kochunov, B.T. Thomas Yeo, Daniel Margulies, Jonathan Smallw, Simon B. Eickhoff, Boris C. Bernhardt

**Affiliations:** Otto Hahn Group Cognitive Neurogenetics, Max Planck Institute for Human Cognitive and Brain Sciences, Leipzig, Germany; INM-7, FZ Jülich, Jülich, Germany; Institute of Systems Neuroscience, HHU Duesseldorf, Duesseldorf, Germany; Center for the Developing Brain, New York City, USA; INM-1, FZ Jülich, Jülich, Germany; Multimodal Imaging and Connectome Analysis Lab, McConnell Brain Imaging Centre, Montreal Neurological Institute and Hospital, McGill University, Montreal, Quebec, Canada; Department of Data Science, Inha University, Incheon, South Korea; 6. University of Baltimore, Baltimore, USA; Department of Psychiatry, Cambridge University, Cambridge UK; Maryland Psychiatric Research Center, University of Maryland School of Medicine, Baltimore, Maryland, US; Department of Electrical and Computer Engineering, National University of Singapore, Singapore, Singapore; Centre for Sleep and Cognition (CSC) & Centre for Translational Magnetic Resonance Research (TMR), National University of Singapore, Singapore, Singapore; N.1 Institute for Health & Institute for Digital Medicine (WisDM), National University of Singapore, Singapore, Singapore; Martinos Center for Biomedical Imaging, Massachusetts General Hospital, Charlestown, Massachusetts, United States of America; Integrative Sciences and Engineering Programme (ISEP), National University of Singapore, Singapore, Singapore; Neuroanatomy and Connectivity Lab, Institut de Cerveau et de la Moelle epiniere, Paris, France; Department of Psychology, Queen’s University, Kingston, Ontario, Canada

## Abstract

Brain structure scaffolds intrinsic function, supporting cognition and ultimately behavioral flexibility. However, it remains unclear how a static, genetically controlled architecture supports flexible cognition and behavior. Here, we synthesize genetic, phylogenetic and cognitive analyses to understand how the macroscale organization of structure-function coupling across the cortex can inform its role in cognition. In humans, structure-function coupling was highest in regions of unimodal cortex and lowest in transmodal cortex, a pattern that was mirrored by a reduced alignment with heritable connectivity profiles. Structure-function uncoupling in non-human primates had a similar spatial distribution, but we observed an increased coupling between structure and function in association regions in macaques relative to humans. Meta-analysis suggested regions with the least genetic control (low heritable correspondence and different across primates) are linked to social cognition and autobiographical memory. Our findings establish the genetic and evolutionary uncoupling of structure and function in different transmodal systems may support the emergence of complex, culturally embedded forms of cognition.

## INTRODUCTION

Cognition helps an animal to satisfy core biological goals in a changing environmental context. In humans, cognition allows our species to successfully navigate through a broad array of situations and socio-cultural contexts. Although the need for flexible cognition is well-established, it remains unclear how a relatively static brain organization can give rise to functional patterns with sufficient flexibility to navigate complex culturally-rich landscapes, such as those found in human societies.

Contemporary perspectives suggest that higher-order, abstract, cognition is grounded in a cortical organization that encompasses parallel axes of microstructural differentiation and function ^1^[for nomenclature: **Supplementary Table 1**]. On the one hand, sensory/motor systems as well as unimodal association cortices are involved in operations related to perceiving and acting in the outside world. These systems are differentiated from transmodal systems that are less tied to a specific modality, and are increasingly engaged in abstract and self-generated cognition together with communication with the “internal milieu” ^1–4^. These functional differences are reflected in well-established differences in the microstructure of sensory/motor and transmodal cortex. Histological studies have shown that unimodal sensory and motor regions show more distinctive lamination patterns relative to agranular/dysgranular transmodal cortex with less apparent lamination ^5^. Complementing these findings, *in vivo* studies have shown that transmodal regions have overall lower myelin content ^6–9^, yet more complex dendritic arborization patterns which could facilitate integrative processing and increased potential for plastic adaptations ^10^. According to its classic definition ^1^, transmodal cortex encompasses both paralimbic cortices as well as heteromodal association networks ^11^, notably the default mode and fronto-parietal functional networks that are specifically expanded in humans^11^. These latter two networks are known to participate in a broad class of abstract cognitive processes, including autobiographical memory ^12, 13^, language ^14–16^, as well as executive control ^2, 17, 18^.

*Post-mortem* studies in non-human animals together with emerging data in humans ^19–21^ suggest that regions with a similar cytoarchitecture are also more likely to be structurally and functional interconnected, an observation framed as the “structural model” of brain connectivity ^22, 23^. Yet, how mappings in cortical structure and function vary across different cortical areas remains to be established. Recent *in vivo* work suggests that structure-function coupling as measured by the association of white matter tractography and functional connectivity is progressively diminished towards transmodal cortex relative to sensory/motor and unimodal cortex ^24–26^. Similar findings can also be observed when studying associations between cortical microstructure based on T1-weighted/T2-weighted (T1w/T2w) and functional connectivity ^21^, collectively pointing to a differential organization of transmodal systems in terms of cortical structure and function relative to unimodal systems ^21, 27, 28^.

Since transmodal systems are assumed to play a role in cognition are less constrained to specific modalities of information, a reduction in the constraining influence of cortical structure on function in transmodal systems may be an important evolutionary adaptation supporting human cognition ^1, 3, 11, 26^. For example, heteromodal regions have been reported to show increased expansion in surface area and untethering from external and internal inputs, processes potentially linked to human cognition ^1, 11^. To better understand these untethered regions of cortex, our study set out to understand whether associations between cortical structure and function may enable an architecture giving rise to abstract human cognition using two related approaches. First, we mapped structure-function relationships across the cortex in humans, and examined whether these profiles were heritable. Heritability serves as a backbone for evolutionary change, as natural selection acts upon inherited traits under variation ^29^. In humans, patterns of cortical microstructure and functional connectivity are heritable, indicating partial genetic control over individual variation ^30–36^. Complementing the heritability assessment in humans, we examined how structure-function associations seen in our species are preserved in non-human primates (NHP). Phylogenetic comparisons between humans and NHP can shine a light into evolutionary conservation and adaptation across primates ^37, 38^. Previous work has shown that spatial variations in cortical microstructure ^14, 39^ and functional connectivity seen in humans ^40, 41^ are already present in NHP, but that there several evolutionary innovations emerging throughout the human lineage. For example, comparing markers of cortical myelin content between NHP and humans, it was shown that the arcuate fasciculus underwent evolutionary modifications particularly in humans ^14^. This finding is paralleled by observations showing marked expansions of transmodal surface area in humans compared to NHP ^39, 42^. These expansions may also be reflected in relocations of heteromodal functional networks when comparing humans and macaques ^40^. However, to what extent microstructure-functional connectivity relationships in NHPs mirror those in humans, remains to be investigated.

By combining heritability and cross-species approaches to probe structure-function associations within a single study, we hope to understand processes that underpin abstract cognition in our species and to identify uniqueness of human structure-function coupling and decoupling in transmodal areas. Structure and function were measured using node-level correlation of T1w/T2w microstructure profile covariance (MPC) ^21^ and resting-state functional connectivity (rsFC). To study heritability, we analyzed the pedigree design and multimodal imaging data of the Human Connectome Project S1200 release ^43^. Equivalent analyses in macaques based on the PRIMate-Data Exchange repository ^37^ examined phylogenetic differences in microstructural and functional organization. We combined node-level network neuroscience approaches with the use of unsupervised dimensionality reduction techniques, which identified large-scale microstructural and functional gradients and provided a coordinate system to map genetic and evolutionary influences on cortical organization ^28, 44, 45^. Finally, we contextualized the likely functional profile of these regions through meta-analytical data from the task-based fMRI literature ^46^. We performed various robustness and replication analyses to assess stability of our findings.

## RESULTS

### Cortex wide decoupling of function and structure (Figure 1)

We first established the spatial distribution of structure-function coupling in the human brain, using node-level association analyses. Specifically, we mapped how patterns of functional connectivity reflect similarity of cortical microstructure across all cortical regions, using the S1200 sample of the Human Connectome Project young adult dataset ^43^ (see *Materials and Methods* for details on participant selection and image processing). To construct microstructure profile covariance (MPC) matrices, we sampled intracortical T1w/T2w values at 12 different cortical depths ^21^, and correlated dept-wise cortical intensity profiles between the parcels (**Figure 1A**). To control for curvature effects, equivolumetric surfaces were used^47^. Resting-state functional connectivity (rsFC) matrices were calculated by cross-correlating the neural time series between all pairs of 400 cortical nodes ^48^ (**Figure 1B**). Correlating node-wise patterns in both measures, averaged across participants (**Figure 1C**), we observed an overall edge-level association between MPC and rsFC in line with the predictions of the structural model ^20^. As expected, however, there was also a progressively decreasing correspondence between MPC and rsFC (r=0.525, p<0.001) along a sensory-fugal gradient of cytoarchitectural classes, capturing cytoarchitectural complexity [^49^, for nomenclature: **Supplementary Table 1** and **Figure 1C**] from high correlations in primary regions (mean±sd: r=0.450 0.181), to weak correlations unimodal association cortices (r=0.134 0.263), followed by close to zero correlations in transmodal cortex (heteromodal association cortices, r=0.049 0.198; paralimbic areas, r=-0.003 0.146). Findings were consistent in a replication sample (N= 50, Royer et al, in prep) and when using an alternative node parcellation (**Supplementary Results**). Relative to sensory/motor and unimodal areas, transmodal systems show the strongest decoupling between cortical microstructure and functional connectivity.

**Figure 1.**
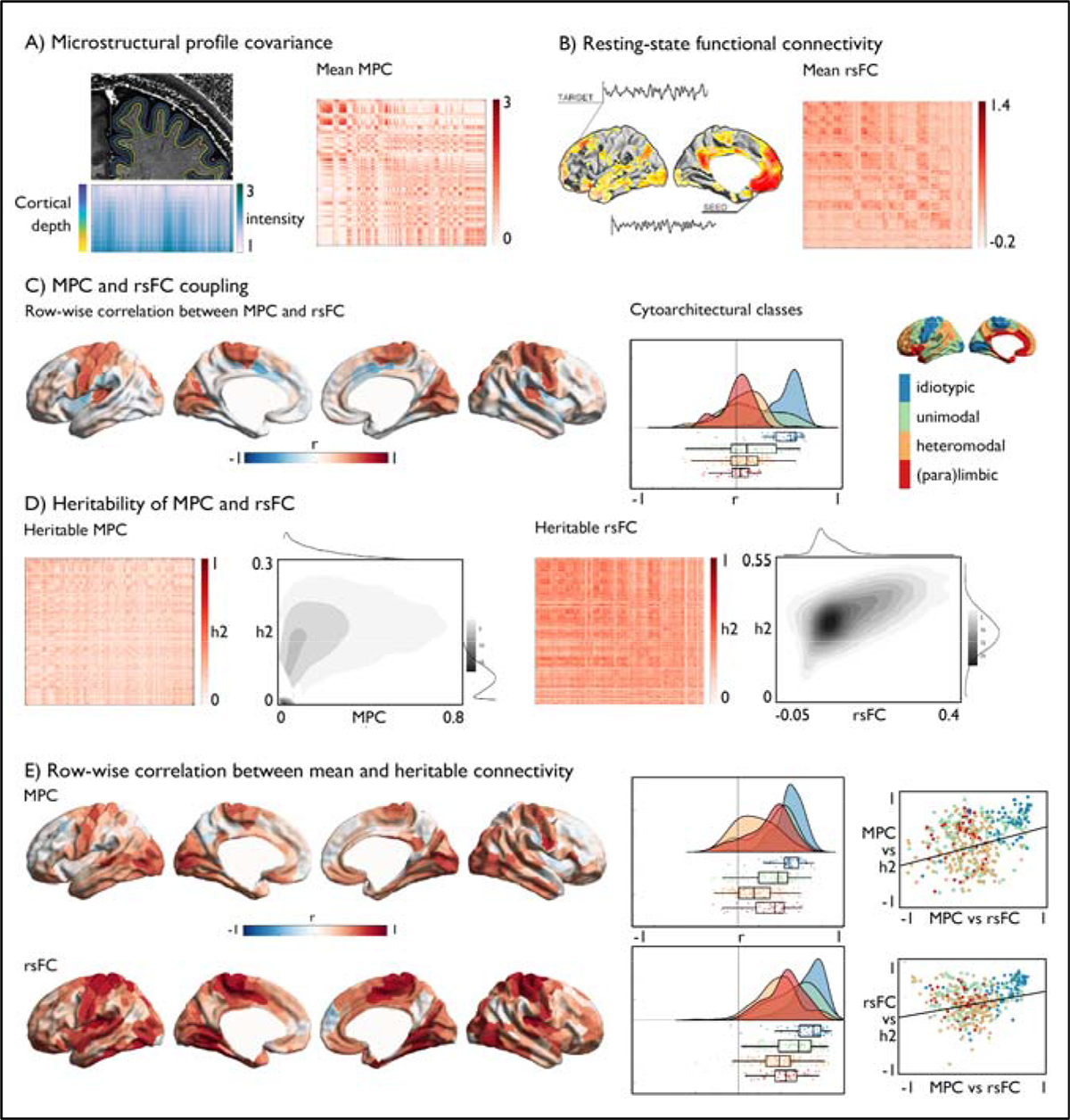
Structure-function coupling and heritability in human cortical regions. **A**) Microstructural profile covariance (MPC) was chosen to map networks of microstructural similarity for each cortical node, sorted along cytoarchitectural class ^21, 49^; **B**) Resting-state functional connectivity (rsFC) analysis maps nodal patterns of intrinsic functional connectivity, sorted along cytoarchitectural class; **C**) Row-wise coupling of MPC and rsFC, as well as correspondence averaged per cytoarchitectural class, *Middle*: raincloud plot of distribution within cytoarchitectural classes; *Right*: Reference visualization cytoarchitectural class; **D**) Heritability of MPC and rsFC and node-wise correspondence between mean and heritable connections; **E**) Row-wise association between mean and heritable seed-wise connectivity – reflecting genetic control over connectivity profiles, *Middle:* distribution per cytoarchitectural class, and *Right* correlation between MPC-rsFC coupling and mean MPC – heritable MPC / rsFC respectively.

### Genetic control over structural and functional connectivity profiles

Having documented reductions in the association between in MPC and rsFC in transmodal cortices, we next examined whether this difference is heritable, i.e., under genetic control. The HCP S1200 sample contains both unrelated as well as genetically related individuals, allowing us to analyze heritability through maximum likelihood analysis. We computed the node-wise heritability of MPC, and then correlated the heritability with mean MPC to index genetic control. A similar approach was carried out for rsFC connectivity profiles. Both analyses were again performed at a whole network and node level. In this context, high correlations suggest that edges with high MPC/rsFC correspondence also show high heritability, whereas low correlations suggest low heritability. At the whole network level, MPC was heritable (mean±sd h2=0.168±0.033), but effects appeared weaker than for rsFC (mean±sd h2=0.340±0.043). Similarly, MPC and its heritability were correlated (rho=0.283), but this correlation was higher for rsFC (Spearmans rho=0.537). At a node level, primary sensory/motor regions showed positive correlation between mean and heritable connectivity profiles for both MPC and rsFC, whereas transmodal regions showed less correlations between mean and heritable connectivity profiles in both measures (MPC: mean±sd primary (idiotypic): r=0.482±0.145; unimodal: 0.318±0.219; heteromodal:0.168±0.191; paralimbic: 0.293±0.206 and rsFC: primary: r=0.641±0.185; unimodal: 0.522±0.204; heteromodal:0.371±0.210; paralimbic: 0.431±0.172). Correlating mean-heritability correspondence within each modality and the correlation between modalities, while controlling for spatial autocorrelations using spin-tests ^50^, we found a moderately positive association for both MPC (r=0.335, p_spin_=0.02) and rsFC (r=0.292, p_spin_=0.02). This indicates that whereas in primary regions the correlation between MPC and rsFC as well as the correspondence of mean and heritable connectivity profiles is strong, there is less consistent correspondence between mean and heritable connectivity patterns in regions showing low structure-function coupling, and particularly in heteromodal areas. In other words, reduced structure-function coupling in human transmodal regions was paralleled by reductions in genetic control over both MPC and rsFC.

*<u>Correspondence between microstructure and function in non-human primates (</u>Figure 2)* Having established that structure-function is reduced in transmodal cortex in humans, we next examined whether the above structure-function associations are also seen in other primates, by evaluating 19 macaques from the PRIMate-Data Exchange who had microstructural MRI and resting-state fMRI available ^37^. Analogous to the human analysis, we created MPCs using multiple equivolumetric surfaces between pial and grey matter/ white matter surfaces to extract depth dependent T1w/T2w profiles in each monkey (details in **Supplementary Results**). We carried out an rsFC analysis, and aligned humans and macaques using a recently introduced cross-species procedure ^40^. Comparing edges of MPC and rsFC in macaques, we found an overall correlation between MPC and rsFC (r=0.22, p<0.0001). Correspondence between MPC and rsFC was similar in both species (r=0.38, p_spin_=0.035), albeit stronger in macaques (t=6.48, p<0.001, CI [0.109 0.204]). Comparing the associations at the microstructural-level, using cytoarchitectural classes in both species (**Supplementary Table 1**), we found no difference between coupling in paralimbic regions (uncoupled in humans and macaques; t=1.252, p>0.1) and primary sensory/motor regions (coupled in humans and macaques; t=-0.33, p>0.1). However, both unimodal (t=6.626, p<0.001) and heteromodal association regions (t=7.070, p<0.001) were generally more coupled in macaques than in humans. This analysis shows that while humans and macaques show broadly similar spatial trends of reduced structure-function correspondence from sensory to transmodal areas, uni- and heteromodal association cortices are characterized by reductions in structure-function coupling in humans compared to NHPs.

**Figure 2.**
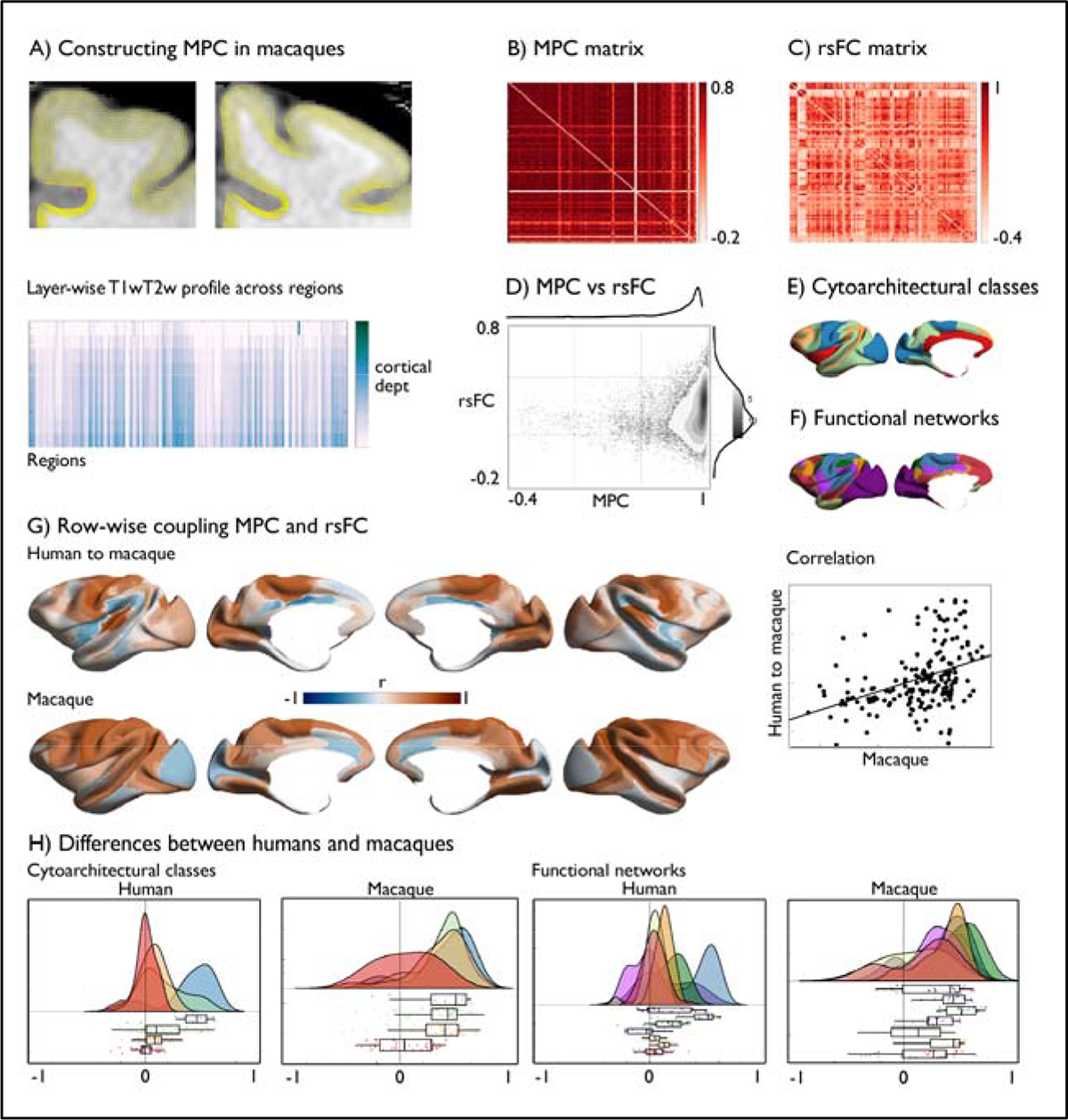
Microstructure-function coupling in macaques. **A**) Creating MPC in macaques; **B**) MPC matrix in macaques, ordered along cytoarchitectural class based on ^49^ and Markov labels ^51^; **C**) rsFC matrix in macaques, ordered along cytoarchitectural class; **D**) Correspondence between MPC and rsFC in macaques; **E**) Cytoarchitectural classes; **F**) Functional communities based on ^27^; **G**) Row-wise association of MPC and rsFC; upper panel: human map in macaque space; lower panel: macaque map; right: scatter between human and macaque MPC-rsFC tethering; **H**) Rainbow plots of difference between humans and macaques as a function of cytoarchitectural class and functional communities ^27^ in macaque space.

### Organizational differences in microstructure and functional connectivity (Figure 3)

To further understand the genetic basis of MPC and rsFC, we evaluated the correspondence in main organizational axes of MPC/rsFC and their node-wise heritability. Applying non-linear dimensionality reduction ^52, 53^, we observed that both MPC and rsFC followed similar spatial axes as their heritability (**Figure 3A**), with principal axes of mean and heritable highly correlated for both MPC (r=0.808, p_spin_<0.0001) and for rsFC (r=0.892, p_spin_<0.0001). Next, we calculated the nodal difference between standardized and aligned MPC and rsFC principal gradients (Δ_MPC-rsFCG1_). In line with previous reports ^21^, gradients were most uncoupled in transmodal regions. The MPC gradient ran from sensory/motor to paralimbic regions, while the rsFC gradient radiated from sensory/motor to heteromodal networks (**Figure 3B**). We then selected homologue gradients of MPC and rsFC in macaques (**Supplementary results**) and again observed the most marked difference within transmodal networks. There was also a moderate positive correlation between the difference between MPC and rsFC organization across species (r=0.33, p_spin_ =0.04). This suggests that the differentiability of topological organization of transmodal regions for MPC and rsFC is present in at least two primate species.

**Figure 3.**
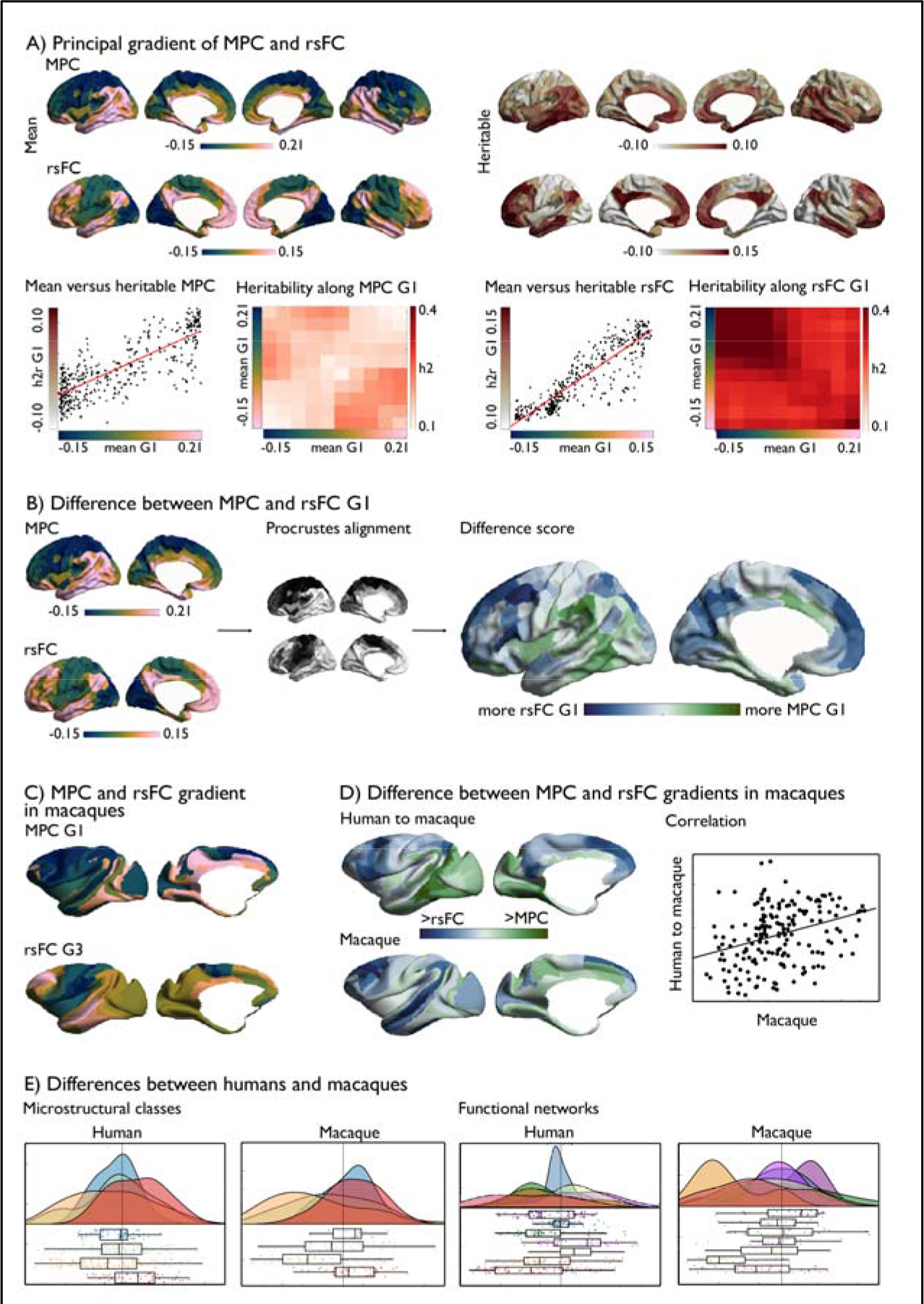
Difference in organizational gradients of MPC and rsFC in humans and macaques. **A)** Principal gradient of MPC and rsFC, *left:* gradients based on mean data on MPC and rsFC and *right* gradients of heritable data alone, *lower left* panel: mean versus heritable MPC G1, as well as heritably along the principal mean gradient in MPC; *lower right* panel: mean versus heritable MPC G1, as well as heritably along the principal mean gradient in MPC; **B**) *Left panel*: Principal MPC and rsFC gradient; *Middle panel*: alignment; *Right panel*: Δ_rsFC-MPCG1_; **C**) Principal gradient of MPC and tertiary gradient of rsFC in macaques; **D**) Difference between principal gradients of MPC and rsFC in humans, mapped to macaque space, and difference between corresponding gradients in macaques (lower panel); *right*: correlation between human and macaque maps; **E**) Raincloud plots of organizational differences as a function of cytoarchitectural class ^49^, and functional networks ^27^ in humans and macaques.

### Multiscale quadrants of structure-function coupling (Figure 4)

Our final analysis examines the most likely functional consequences of structure-function decoupling for human cognition. The axes of structure-function coupling and gradient differences offer a two-dimensional coordinate system to visualize and conceptualize how this cortical organization is linked to function. As expected, the axis describing structure-function coupling differentiates primary regions from transmodal regions (upper and lower half of the quadrant). Conversely the MPC-rsFC gradient difference axis differentiates heteromodal from paralimbic regions. To underscore the ability of this space to represent cortical organization, we assessed its ability to differentiate different motifs of neural structure, evolution, and function (See *Methods*).

We first projected cytoarchitectural classes and intrinsic functional communities into our two-dimensional coordinate system to understand the large-scale communities linked to reduced structural-function coupling (**Figure 4A**). In humans and macaques, both upper quadrants include sensory/motor and unimodal networks. However, while in macaques the left upper quadrant is also occupied by fronto-parietal and default mode networks, these networks are in the lower left quadrant in humans. The lower right quadrant is dominated by limbic and ventral attention network in humans, and limbic networks in macaques. Comparing the relative number of nodes distributed in each quadrant, more areas were uncoupled in humans relative to macaques (χ^2^: 14.57, p=0.0013), particularly in the left half of the quadrant (χ^2^: 10.24, p=0.0014).

**Figure 4.**
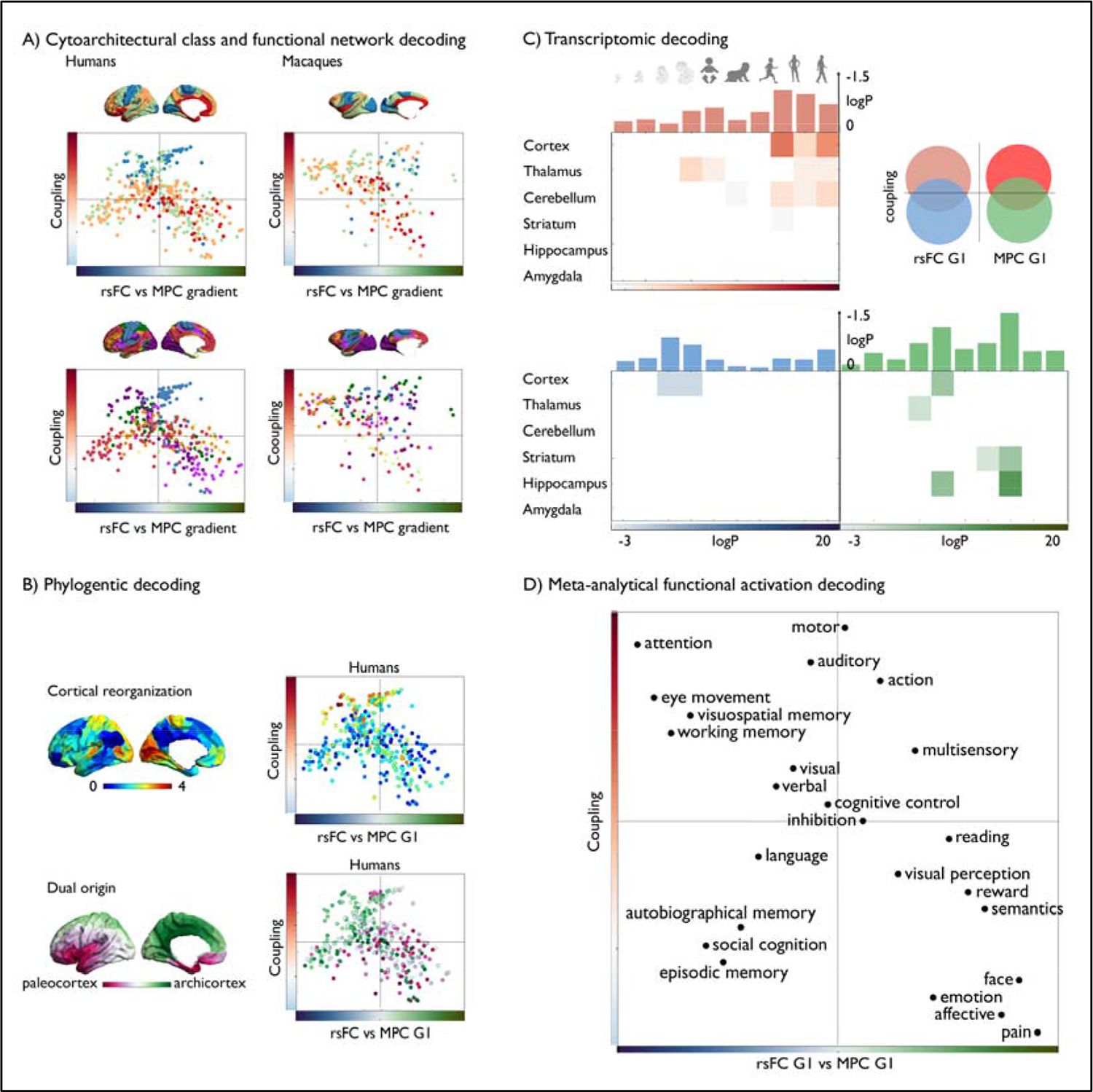
Multiscale quadrants of structure-function coupling; **A**) Cytoarchitectural class ^49^ and functional community ^27^ decoding along 2D model of the difference between microstructural and functional connectivity gradients in humans and macaques (x-axis) and microstructure-functional connectivity coupling (y-axes); **B**) Phylogenetic models using cortical reorganization ^40^ and a model of dual patterning in the cerebral cortex ^54^**; C**). Transcriptomic developmental decoding of coupling and gradient differences between structure and function, Red/Blue/Green tones represent the log transformed false discovery rate (FDR)-corrected p-values [-20-3]. The bar plot above represents the log transformed (FDR)-corrected p-values, averaged across all brain structures. Red indicates the genes that were attributed to the left upper quadrant, blue indicates the values were higher for the functional end of the difference gradient and untethered, whereas green reflects the microstructural apex and untethered. Only values that are below FDRq<0.05 are displayed; **D**) 2D projection of NeuroSynth meta-analysis of regions of interest along ⊗_rsFC-MPCG1_ map (x-axis) and structure-function uncoupling (y-axis) using 24 topic terms.

In line with the observed cross-species differences, the quadrants furthermore reflected a differentiation of cortical reorganization between humans and macaques reported in prior findings ^40^, and suggested different archi-vs paleocortical trends relative to the dual origin model of cortical differentiation ^54^ (**Figure 4B, Supplementary Results**). Different biological pathways were furthermore suggested by projecting gene expression maps from the Allen Human Brain Atlas (AHBA) into the 2D space ^55^. Indeed, the lower left quadrant associated with genes mainly expressed in prenatal states in cortical areas, while the lower right quadrant was additionally associated with prenatal expression of nodes in the hippocampus together with the cortical regions and thalamus, striatum and hippocampus postnatally (**Figure 4C, Supplementary Results**). Thus, the genetic uncoupling observed in twins in our main analyses may reflect differential time-windows of developmental expression in cortical and non-cortical regions.

Finally, we identified the most likely cognitive consequences of the structure-function decoupling in transmodal cortex. To this end we mapped cognitive ontologies based on NeuroSynth into the two-dimensional space ^46^ (**Figure 4D**). Quadrants with high structure-function coupling included primary and unimodal regions and related to terms such as “working memory”, “attention”, and “executive control” (left upper quadrant) and “action” as well as “multisensory processes” (right upper quadrant). Quadrants with low coupling were made of the paralimbic regions and related to affective processes (“emotion”, “reward”, “pain”; lower right) and heteromodal regions (“episodic memory”, “social cognition”, lower left). This indicates that different quadrants also reflected different behavioral and cognitive processes.

## DISCUSSION

Our study set out to understand how flexible forms of cognition and behavior emerge from the cortex, motivated by an emerging hypothesis linked to reduced grip of cortical structure on function in transmodal systems, possibly enabling functional processes are less constrained to specific modalities of information ^1,11, 20, 21, 23, 24, 28, 56^. Our analysis confirmed prior observations that human brain structure and function are generally least coupled in transmodal cortex. This decoupling mainly related to a reduction in genetic control onto both the structure and function of heteromodal but not paralimbic components of transmodal cortex, including default mode and fronto-parietal functional networks. Analyses in non-human primates also showed a decoupling of structure and function may be an architecture broadly conserved across primate species ^57^. However, the decoupling observed in macaques was less pronounced than in humans, particularly in heteromodal but again not paralimbic transmodal regions. Heteromodal and paralimbic regions were found to reflect distinctive functional attributes, specifically showing an implication of heteromodal regions in social cognition and autobiographical memory, while paralimbic regions were more closely implicated in affective/motivational processes. Both these functional types were furthermore found to relate to different gene expression pathways, indicating different biological origin. Collectively, our systematic assessment of genetic and evolutionary factors contributing to cortical structure-function associations help to disentangle macroscale cortical systems, particularly the different components of transmodal cortex that are believed to be involved in culturally enriched thought that is at the core of human uniqueness.

There are a number of reasons to suspect that regions where MPC and rsFC diverge the most may support heightened experience-induced development and plasticity ^58^. For example, transitions from late childhood to early adulthood are reflected in altered structure-function coupling ^26^ and these consistent changes in cortical microstructure and myelination patterns are seen in transmodal regions ^59^. A protracted and reduced myelination of axons may aid the coordination of distributed neural activity to occur later in life, allowing more flexible neural motifs to emerge over time ^26, 60^. Moreover, regions of transmodal cortex are associated with higher degrees of synapse formation and growth, as measured by aerobic glycolysis in the adult brain^61^. The apparent reduction in the genetic control associated with reduced structure-function decoupling in transmodal cortex established by our study may facilitate these forms of cortical adaptation that continue to occur after birth. Consistent with this view, regions within the default mode and fronto-parietal networks show most evidence of dynamic changes and experience-dependent plasticity at both short and longer time scales ^62–64^. For example, studies of spontaneous cognition suggest that dynamic reconfigurations of default mode and fronto-parietal systems are linked to dynamic changes in neural activity that are correlated with patterns of ongoing thought ^65, 66^.

Our functional analysis is also consistent with the view that structure-function decoupling is linked to forms of cognition informed through experience. We found that regions with the strongest structure function coupling are linked to cognitive terms such as ‘attention’ and ‘working memory’. Conversely, uncoupled regions that were similar to humans and macaques were linked to emotional and motivational states, processes that are important in most if not all mammals ^1^. Notably, however, regions in which structure-function decoupling was most unique to humans were associated with functions such as “social-cognition” and “autobiographical memory”. The emergence of declarative memory, and explicit social functions have been reported in non-human primates ^67, 68^ but, at the same time, are thought to be foundational processes upon which cultural learning in our species are grounded. More generally, our analysis provides important insight into the broader organization framework of human cognition. The microstructural gradient captures spatial shifts in cortical lamination patterns ^21^ and represents a ‘sensory-fugal’ axis that runs towards paralimbic cortices ^1^, implicated in predictive processing of allostatic needs and motivation ^69^. Also anchored on sensory systems but radiating towards heteromodal networks such as the default mode network, the principal functional gradient closely relates the distance of brain regions from sensory input ^3, 70, 71^ and may capture part of a divergent pattern of connectivity that protrudes from the sensory-fugal hierarchy. Here, heteromodal regions show most long-range functional connectivity patterns, linked to changes in supra-granular layers ^72–74^. As such, this axis ranging from sensory to heteromodal regions may be linked to the distributed processing of information that is relatively uncoupled from sensory input, and optimally positioned for the integration of external and internal information needed for experience informed modes of thought such social cognition or autobiographic memory processes ^3, 11^.

Our data is also consistent with the notion that rapid evolutionary expansion of the cerebral cortex may have shifted away from bottom-up activity cascades in primary sensory-motor regions towards a network of relatively untethered, long-distance, and increasingly parallelized heteromodal regions ^11, 74^. We observed a relationship between the uncoupling of structure and function in transmodal regions and maps of functional reorganization between macaques and humans ^40^ and the dual origin theory of cortical organization ^54, 75^. The latter argues that neural differentiation progressively stems from an archicortical and paleocortical origin ^54^, and our findings suggest that archicortical trends preferentially relate to heteromodal, while paleocortical trends encompass paralimbic regions. Moreover, the closer regions were to either origin, the stronger their structure-function uncoupling. Together these observations suggest that differentiable phylogenetic processes may underlie transmodal uncoupling in heteromodal and paralimbic cortices.

In line with potentially different implicated biological pathways shaping transmodal regions, we observed different transcriptomic associations across the sensory-fugal axis more generally and heteromodal and paralimbic regions specifically ^20, 23, 56^. Coupled regions showed a close association with postnatal expression of genes in neocortical regions, thalamus, and cerebellum, in line with cerebellar-thalamic-cortical circuits ^76^. Conversely, we found that paralimbic transmodal regions were associated with prenatal genetic expression in the neocortical regions, thalamus, and hippocampus and postnatal genetic expression in striatum and hippocampus, whereas heteromodal regions uniquely related to genes expressed in neocortical regions in prenatal stages. The different expression time-windows observed may be a genetic indicator of how heteromodal regions, associated with genes expressed early in development, can develop more rich connectivity profiles along the “older get richer” principle ^11, 77^. Conversely those regions associated with genetic expression later in development are under heightened genetic control, and not only include primary regions that show a tight coupling of structure and function, but also paralimbic regions, linked to the striatum. Thus, transmodal decoupling may be understood from a perspective reaching beyond the cortex, guided by sub-cortex such as the thalamus and striatum ^76, 78, 79^. In particular, thalamus, as well as both the cerebellar hemispheres, show recent evolutionary alterations ^80, 81^ and show increased connectivity to the granular (dorsal) prefrontal cortices ^82, 83^. As such it is possible that the uncoupling of particularly heteromodal regions observed in humans may build upon evolutionary alterations not only in the cortex but also sub-cortex, ultimately supporting associated key features of human cognition.

In conclusion, our study set out to understand how flexible cognition emerges from a static genetically controlled neural architecture. Our analyses suggest that the previously documented pattern of reduced structure function coupling in transmodal regions of cortex is heritable in humans and broadly conserved in different primate species. Importantly, in non-human primates this pattern was reduced relative to humans, and a functional meta analyses indicated that this difference was most likely associated with social cognition and autobiographical memory. Together these data are consistent with the hypothesis that during evolution brain organization has been shaped to support structure-function decoupling to facilitate types of cognition that can be enriched through learning and social interactions, enabling important features of cultural processes in our species.

## Materials and methods

### HCP sample

#### Participants and study design

We used publicly available data from the Human Connectome Project S1200 release (HCP; http://www.humanconnectome.org/), which comprised data from 1206 individuals (656 females) that are made up by 298 MZ twins, 188 DZ twins, and 720 singletons, with mean±SD age 28.8±3.7 years (range = 22–37 years). We included individuals for whom the scans and data had been released after passing the HCP quality control and assurance standards. The full set of inclusion and exclusion criteria are described elsewhere ^7, 43^. In short, the primary participant pool comes from healthy individuals born in Missouri to families that include twins, based on data from the Missouri Department of Health and Senior Services Bureau of Vital Records. Additional recruiting efforts were used to ensure participants broadly reflect ethnic and racial composition of the U.S. population. Healthy is broadly defined, in order to gain a sample generally representative of the population at large. Sibships with individuals having severe neurodevelopmental disorders (e.g., autism), documented neuropsychiatric disorders (e.g. schizophrenia or depression) or neurologic disorders (e.g. Parkinson’s disease) are excluded, as well as individuals with diabetes or high blood pressure. Twins born prior 34 weeks of gestation and non-twins born prior 37 weeks of gestation are excluded as well. After removing individuals with missing structural and functional imaging data our sample consisted of 992 (529 females) individuals (including 255 MZ-twins and 150 DZ-twins) with a mean age of 28.71 years (SD =3.72, range =22-37).

#### Structural imaging processing

MRI protocols of the HCP are previously described ^7, 43^. In short, MRI data used in the study were acquired on the HCP’s custom 3T Siemens Skyra equipped with a 32-channel head coil. Two T1w images with identical parameters were acquired using a 3D-MPRAGE sequence (0.7 mm isotropic voxels, matrix = 320 × 320, 256 sagittal slices; TR = 2,400 ms, TE = 2.14 ms, TI = 1,000 ms, flip angle = 8°; iPAT = 2). Two T2w images were acquired using a 3D T2-SPACE sequence with identical geometry (TR = 3,200 ms, TE = 565 ms, variable flip angle; iPAT = 2). T1w and T2w scans were acquired on the same day. The pipeline used to obtain the Freesurfer-segmentation is described in detail in a previous article ^7^ and is recommended for the HCP-data. The pre-processing steps included co-registration of T1- and T2-weighted scans, B1 (bias field) correction, and segmentation and surface reconstruction using FreeSurfer version 5.3-HCP. Using these data, the equidistant surfaces are computed for MPC measurement.

#### Parcellation approach

For our main analysis, we used a parcellation scheme ^48^ that combines local gradient and global similarity approaches via gradient-weighted Markov Random models. The parcellation has been evaluated with regards to stability and convergence with histological mapping and alternative parcellations. Here, we focused on the granularity of 400 parcels. We additionally evaluated results using the Glasser 360 atlas ^84^.

### Cortical microstructure and microstructural covariance networks

We estimated MPC using myelin-sensitive MRI, in line with the previously reported protocol ^21^, in the S1200 HCP sample. The myelin-sensitive contrast was T1w/T2w from the HCP minimal processing pipeline, which uses the T2w to correct for inhomogeneities in the T1w image. We generated 12 equivolumetric surfaces between the outer and inner cortical surfaces. The equivolumetric model compensates for cortical folding by varying the Euclidean distance ρ between pairs of intracortical surfaces throughout the cortex to preserve the fractional volume between surfaces. ρ was calculated as follows for each surface (1):

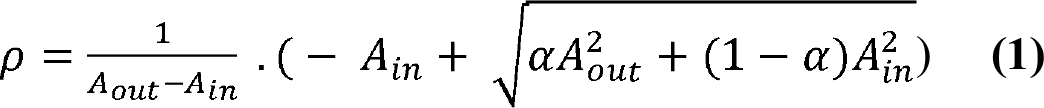

in which α represents a fraction of the total volume of the segment accounted for by the surface, while A_out_ and A_in_ represents the surface area of the outer and inner cortical surfaces, respectively. We systematically sampled T1w/T2w values along 64,984 linked vertices from the outer to the inner surface across the whole cortex. Subsequently, we computed the average value of T1w/T2 in each of the 400 parcels of the Schaefer atlas ^48^. In turn, MPC *(i*,*j)* for a given pair of parcels *i* and *j* is defined by (2):

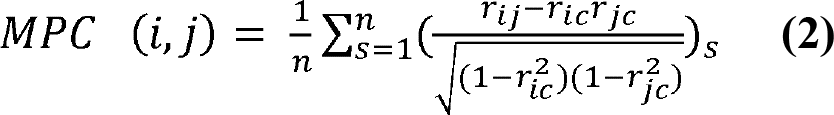

in which *s* is a participant and *n* is the number of participants, *r_ic_* the correlation coefficient of the intensity profile at node *i* with the average intensity profile across the entire cortex, and *r_jc_* the correlation of the intensity profile at node *j* with the average intensity profile across the cortex. We used the MPC for further analysis.

#### Functional connectivity

Functional connectivity matrices were based on 1 hour of resting-state fMRI data acquired through the HCP ^7^ and made publicly available for download on ConnectomeDB. Functional resting-state MRI data underwent HCP’s minimal preprocessing ^7, 84^. Briefly, for each individual, a functional connectivity matrix was calculated using the correlation coefficient across four minimally preprocessed, spatially normalized, and concatenated to 4 15-min resting-state fMRI scans and co-registered using MSMAll to template HCP 32k_LR surface space ^43^. 32k_LR surface space consists of 32,492 total nodes per hemisphere (59,412 excluding the medial wall). Following average time-series were extracted in each of the 400 cortical parcels ^48^ and individual functional connectivity matrices were computed. The individual functional connectomes were generated by averaging preprocessed timeseries within nodes, correlating nodal timeseries and converting them to z scores. Here we used the individual timeseries of individuals with complete data in the S1200 sample.

### Measuring structure – function coupling

To measure structure-function coupling we performed row-wise correlation along the mean MPC and rsFC connectomes. A similar approach was take to measure genetic coupling. In short, the values of each row in two matrices were selected and correlated and the r-value resulting from this analysis was projected on the surface.

#### Heritability analysis

To investigate the heritability of MPC and intrinsic functional connectomes, we analyzed edge-wise connectomes of both measures in a twin-based genetic correlation analysis. The quantitative genetic analyses were conducted using Sequential Oligogenic Linkage Analysis Routines (SOLAR) ^85^. SOLAR uses maximum likelihood variance-decomposition methods to determine the relative importance of familial and environmental influences on a phenotype by modeling the covariance among family members as a function of genetic proximity. We used a G+E model to assess heritability in the HCP dataset based on prior work indicating G+E is more parsimonious and leads to more reproducible results in this sample ^86^. This approach can handle pedigrees of arbitrary size and complexity and thus, is optimally efficient with regard to extracting maximal genetic information. To ensure that our functional connectivity and microstructural profile covariance measures were conform to the assumptions of normality, an inverse normal transformation was applied.

Heritability (*h^2^*) represents the portion of the phenotypic variance (σ^2^_p_) accounted for by the total additive genetic variance (σ^2^_g_), i.e., *h^2^* = σ^2^_g_/σ^2^_p_. Phenotypes exhibiting stronger covariances between genetically more similar individuals than between genetically less similar individuals have higher heritability. Within SOLAR, this is assessed by contrasting the observed covariance matrices for a neuroimaging measure with the structure of the covariance matrix predicted by kinship. Heritability analyses were conducted with simultaneous estimation for the effects of potential covariates. For this study, we included covariates including age, and sex, age^2^, and age * sex.

### Macaque data

All datasets in this study were from openly available sources. The macaque data stemmed from one cohort (University of California, Davis) of the recently established PRIME-DE (http://fcon_1000.projects.nitrc.org/indi/indiPRIME.html) (<u>Milham et al., 2018</u>). The full data set consisted of 19 rhesus macaque monkeys (*macaca mulatta,* all female, age=20.38 ± 0.93 years, weight=9.70 ± 1.58 kg) scanned on a Siemens Skyra 3T with 4-channel clamshell coil. All the animals were scanned under anesthesia. In brief, the macaques were sedated with injection of ketamine (10 mg/kg), dexmedetomidine (0.01 mg/kg), and buprenorphine (0.01 mg/kg). The anesthesia was maintained with isoflurane at 1-2%. The details of the scan and anesthesia protocol can be found at (http://fcon_1000.projects.nitrc.org/indi/PRIME/ucdavis.html). The neuroimaging experiments and associated procedures were performed at the California National Primate Research Center (CNPRC) under protocols approved by the University of California, Davis Institutional Animal Care and Use Committee ^87^. The resting-state fMRI data were collected with 1.4 × 1.4 × 1.4 mm resolution, TR=1.6s, 6.67 min (250 volumes) under anesthesia. No contrast-agent was used during the scans. Structural data (T1w and T2w) were acquired with 0.3×0.3×0.3 mm resolution (T1w: TR=2500ms, TE=3.65ms, TI=1100ms, flip angle=7 degree, FOV=154mm; T2w: TR=3000ms, TE=307ms).

<u>MRI data processing</u>: The structural T1w and T2w images were preprocessed using the customized HCP-like macaque pipeline (doi:10.5281/zenodo.3888969). Briefly, the preprocessing includes 1) spatial denoising by a non-local mean filtering operation ^88^, 2) brain extraction using ANTs registration with a reference brain mask followed by manually editing to fix the incorrect volume (ITK-SNAP, www.itksnap.org) ^89^; 3) tissue segmentation using ANTs joint label fusion algorithm and surface reconstruction (FreeSurfer)^90^; 4) T1w and T2w alignment (linear) followed by the linear and nonlinear registration to the high resolution template space (0.3mm); 5) the native white matter and pial surfaces were registered to the Yerkes19 macaque surface template ^91^ (Autio et al. 2020).

MPC was computed similarly to the human approach, however due to the thinner cortex in macaques relative to humans we decided to only focus on 9 equidistant surfaces. The macaque monkey intrinsic functional data were preprocessed as described previously in using a customized Connectome Computational System pipeline for nonhuman pipeline ^40^. Briefly, the rsfMRI data were preprocessed including temporal despiking, motion correction, 4D global scaling, nuisance regression using white matter (WM), and cerebrospinal fluid (CSF) signal and Friston-24 parameter models, bandpass filtering (0.01–0.1 Hz), detrending and co-registration to the native anatomical space. The data were then projected to the native mid-cortical surface and smoothed along the surface with FHWM=3mm. Finally, the preprocessed data were down-sampled to a standard 10k (10,242 vertices) resolution surface^91^. Similar with human preprocessing, functional timeseries were averaged within the Markov parcellation ^51^, and a connectivity matrix was constructed.

*Alignment of human to macaque space:* To evaluate the similarity between human and macaque cortical patterns we transformed the human pattern (rsFC-MPC coupling / gradients) to macaque cortex based on a functional-alignment techniques recently developed. This method leverages advances in representing functional organization in high-dimensional common space and provides a transformation between human and macaque cortices ^40^. This, enabled us to directly compare between species within the same space.

#### Gradient decomposition

To compute macroscale gradients, we performed several analysis steps. The input of the analysis was the MPC and rsFC matrix, cut-off at 90% similar to previous studies ^3,21^. To study the relationships between cortical regions in terms of their features, we used a normalized angle similarity kernel resulting in a non-negative square symmetric affinity matrix. Following we used diffusion mapping, a non-linear dimensionality reduction method^52^. The algorithm estimates a low-dimensional embedding from a high-dimensional affinity matrix. In this space, cortical nodes that are strongly interconnected by either many supra-threshold edges or few very strong edges are closer together, whereas nodes with little or no covariance are farther apart. The name of this approach, which belongs to the family of graph Laplacians, derives from the equivalence of the Euclidean distance between points in the embedded space and the diffusion distance between probability distributions centered at those points. It is controlled by a two parameters α and t, where α controls the influence of the density of sampling points on the manifold (α = 0, maximal influence; α = 1, no influence), and t controls the scale of eigenvalues. Based on previous work ^3, 21^ we followed recommendations and set α = 0.5 and t=0, a choice that retains the global relations between data points in the embedded space and has been suggested to be relatively robust to noise. A similar approach was taken for the macaque connectomes.

### Functional decoding

For functional decoding, we selected 24 behavioral paradigms as previously reported in ^3, 21^, averaged in the 400 Schaefer parcels. To perform functional decoding, we averaged the z-scores of NeuroSynth topic terms along the pattern of interest across 20 equally sizes bins and performed weighted averaging, and ranked the values accordingly to capture which functions are associated with MPC-rsFC uncoupling and the difference between MPC and rsFC gradients. Following the ranking of the tasks is projected in 2D space.

### Comparisons between gradients and modalities

To assess the significance of correlations between spatial maps, we used spin-tests to control for spatial autocorrelation when possible ^50^. Difference between two distributions were assessed using a statistical energy test, a non-parametric statistic for two sample comparisons^92^ (https://github.com/brian-lau/multdist/blob/master/minentest.m) and statistical significance was assessed with permutation tests (1000).

#### Phylogenetic maps of cortical reorganization and archi-paleocortex distance

To perform phylogenetic decoding, we used cortical reorganization between macaque monkeys and humans ^40^ (https://github.com/tingsterx/alignment_macaque-human) as well as a model of the dual origin, similar to previous work ^93^. Here we combined the distance to paleo- and archi-cortex in one map, assigning each parcel with the distance closest to either origin.

#### Transcriptomic association analysis

Given the association between phylogeny and ontogeny ^20^, we correlated both maps with post-mortem gene expression data from the Allen Human Brain Atlas (AHBA) ^55^ and evaluated the spatiotemporal time windows in which these genes are most frequently expressed using developmental gene set enrichment analysis ^94^. We assessed spatial correlations of structure-function coupling and the difference between large-scale principal gradients of MPC and rsFC and gene expression patterns. First, we correlated the t-statistics map of the two axes with the post-mortem gene expression maps provided by Allen Institute for Brain Sciences (AIBS) using the Neurovault gene decoding tool ^55, 95^. Neurovault implements mixed-effect analysis to estimate associations between the input map and the genes of AIBS donor brains yielding the gene symbols associated with the input map. Gene symbols that passed a significance level of FDR-corrected p < 0.05 were further tested whether they are consistently expressed across the donors using abagen (https://github.com/rmarkello/abagen), which implements prior recommendations for imaging-transcriptomics studies ^96^. For each gene, we estimated the whole-brain expression map and correlated it between all pair of different donors. Only genes showing consistent whole-brain expression pattern across donors (r>0.5) were retained. In a second stage, gene lists that were significant were fed into enrichment analysis, which involved comparison against developmental expression profiles from the BrainSpan dataset (http://www.brainspan.org) using the cell-type specific expression analysis (CSEA) developmental expression tool (http://genetics.wustl.edu/jdlab/csea-tool-2) ^94^. As the AIBS repository is composed of adult post-mortem datasets, it should be noted that the associated gene symbols represent indirect associations with the developmental data.

### Replication dataset

#### Participants

Data were collected in a sample of 50 healthy volunteers (21 women; 29.82±5.73 years; 47 right-handed) between April 2018 and September 2020 (Royer, in prep). Each participant underwent a single testing session. All participants denied a history of neurological and psychiatric illness. The Ethics Committee of the Montreal Neurological Institute and Hospital approved the study (2018-3469). Written informed consent, including a statement for openly sharing all data in anonymized form, was obtained from all participants.

#### MRI data acquisition

Scans were completed at the Brain Imaging Centre of the Montreal Neurological Institute and Hospital on a 3T Siemens Magnetom Prisma-Fit equipped with a 64-channel head coil. Participants underwent a T1-weighted (T1w) structural scan, followed by resting-state functional MRI (rs-fMRI). In addition, a pair of spin-echo images was acquired for distortion correction of individual rs-fMRI scans. A second T1w scan was then acquired, followed by qT1 mapping. Total scan time for these acquisitions was approximately 45 minutes.

Two T1w scans with identical parameters were acquired with a 3D magnetization-prepared rapid gradient-echo sequence (MP-RAGE; 0.8mm isotropic voxels, matrix=320×320, 224 sagittal slices, TR=2300ms, TE=3.14ms, TI=900ms, flip angle=9°, iPAT=2, partial Fourier=6/8). Both T1w scans were visually inspected to ensure minimal head motion before they were submitted to further processing. qT1 relaxometry data were acquired using a 3D-MP2RAGE sequence (0.8mm isotropic voxels, 240 sagittal slices, TR=5000ms, TE=2.9ms, TI 1=940ms, T1 2=2830ms, flip angle 1=4°, flip angle 2=5°, iPAT=3, bandwidth=270 Hz/px, echo spacing=7.2ms, partial Fourier=6/8). We combined two inversion images for qT1 mapping in order to minimise sensitivity to B1 inhomogeneities and optimize intra- and intersubject reliability ^97, 98^. One 7 min rs-fMRI scan was acquired using multiband accelerated 2D-BOLD echo-planar imaging (3mm isotropic voxels, TR=600ms, TE=30ms, flip angle=52°, FOV=240×240mm^2^, slice thickness=3mm, mb factor=6, echo spacing=0.54ms). Participants were instructed to keep their eyes open, look at a fixation cross, and not fall asleep. We also include two spin-echo images with reverse phase encoding for distortion correction of the rs-fMRI scans (3mm isotropic voxels, TR=4029 ms, TE=48ms, flip angle=90°, FOV=240×240mm^2^, slice thickness=3mm, echo spacing=0.54 ms, phase encoding=AP/PA, bandwidth= 2084 Hz/Px).

#### MRI data pre-processing

Raw DICOMS were sorted by sequence into distinct directories using custom scripts. Sorted files were converted to NIfTI format using dcm2niix (v1.0.20200427; https://github.com/rordenlab/dcm2niix) ^99^, renamed, and assigned to their respective subject-specific directories according to BIDS standards ^100^. Agreement between the resulting data structure and BIDS standards was ascertained using the BIDS-validator (v1.5.10; DOI: 10.5281/zenodo.3762221). All further processing was performed via micapipe, an openly accessible processing pipeline for multimodal MRI data (https://micapipe.readthedocs.io/).

### Data availability

This study followed the institutional review board guidelines of corresponding institutions. All human data analyzed in this manuscript were obtained from the open-access HCP young adult sample (HCP; http://www.humanconnectome.org/). The raw data may not be shared by third parties due to ethics requirements, but can be downloaded directly via the above weblink. Macaque data was obtained from PRIME-DE (http://fcon_1000.projects.nitrc.org/indi/indiPRIME.html; University of California, Davis).

Heritability analyses were performed using Solar Eclipse 8.4.0 (http://www.solar-eclipse-genetics.org), and data on the pedigree analysis is available here: https://www.nitrc.org/projects/se_linux/ ^85, 101^. Gradient mapping analyses was based on BrainSpace (https://brainspace.readthedocs.io/en/latest/). Analysis scripts and additional data are available at (https://github.com/CNG-LAB/cngopen/transmodal_uncoupling/).

## Acknowledgements

We would like to thank the various contributors to the open access databases that our data was downloaded from.

## Author contributions

SLV, SBE and BCB conceptualized the work; SLV, SBE, BCB gave input on analysis; SLV performed the main analysis; TX performed microstructural profile analysis on the macaque data and computed macaque-human gradient correspondence; PK contributed to genetic correlation analysis of the twin model; CP and BCB provided the methods of the microstructural profile covariance; All authors contributed to drafting and revising the manuscript.

## SUPPLEMENTARY RESULTS

### Replication and robustness analysis (Supplementary Figure 1 and 2)

We performed replication analysis in the MICS sample (Royer, in prep). Overall, we found highly convergent patterns (coupling MPC-FC: r=0.736, MPCG1-rsG1: r=0.615). Notably, we observed a difference within the paralimbic network, with the ‘ventral attention’ network being more differentiated from the ‘limbic’ network as defined in ^27^. Also, when analyzing the HCP data using the Glasser parcellation scheme we observed similar dissociations between heteromodal and paralimbic regions as observed using the Schaefer 400 parcellation.

### Variance of principle gradient of MPC and rsFC (Supplementary Figure 3)

We studied whether also the variance (standard deviation) of both measures followed a similar organizational pattern. Indeed, also here we found a close association between the mean gradient pattern and its variance (MPC = 0.98, rsFC = 0.76), indicating that also the individual variation varied along these organizational patterns.

### Macaque gradient validation (Figure 4)

The principal gradient of MPC in macaques showed a sensory-fugal axis in organization of T1wT2w profiles with an apex in superior frontal and sensory-motor regions on the one hand, and an apex in posterior cingulate and inferior temporal lobe on the other hand. The principal gradient of macaques and its human correspondence had a positive correlation (r=0.24, p_spin<0.05). Follow-up analysis indicated this gradient correlated positive with profile skew (r=0.88, p<0.001), rather than the mean regional T1wT2 (r=0.13, p>0.05) (**Supplementary Figure 4**), similar to previous work ^59^. Second, based on previous work (Xu, in prep) we selected the third gradient of rsFC in macaque monkeys to reflect the principal gradient of rsFC in humans. In the current approach, based on averages in the Markov parcellation ^51^, we also observed a positive correlation between the human and macaque gradients (r=0.178, p<0.02).

### Phylogenetic and transcriptomic decoding of the two-dimensional framework of structure-function associations

Structure-function coupling (upper versus lower quadrants) related to cortical re-organization between humans and macaques (r=0.40, p_spin_=0.001) ^40^. The differentiation between large-scale organization axes (left versus right quadrants) governing MPC and rsFC related to distance from archi- and paleocortices (r=0.313, p=0.0001), but not cortical re-organization (r=-0.12, p>0.1).

Moreover, studying the association between spatio-temporal distribution of genetic expression in relation to structure-function mapping we found that the coupling of MPC and rsFC was characterized by increased gene expression in the cortex, thalamus and cerebellum along development (r=0.769, p<0.01).

## SUPPLEMENTARY FIGURES

**Supplementary Figure 1.**
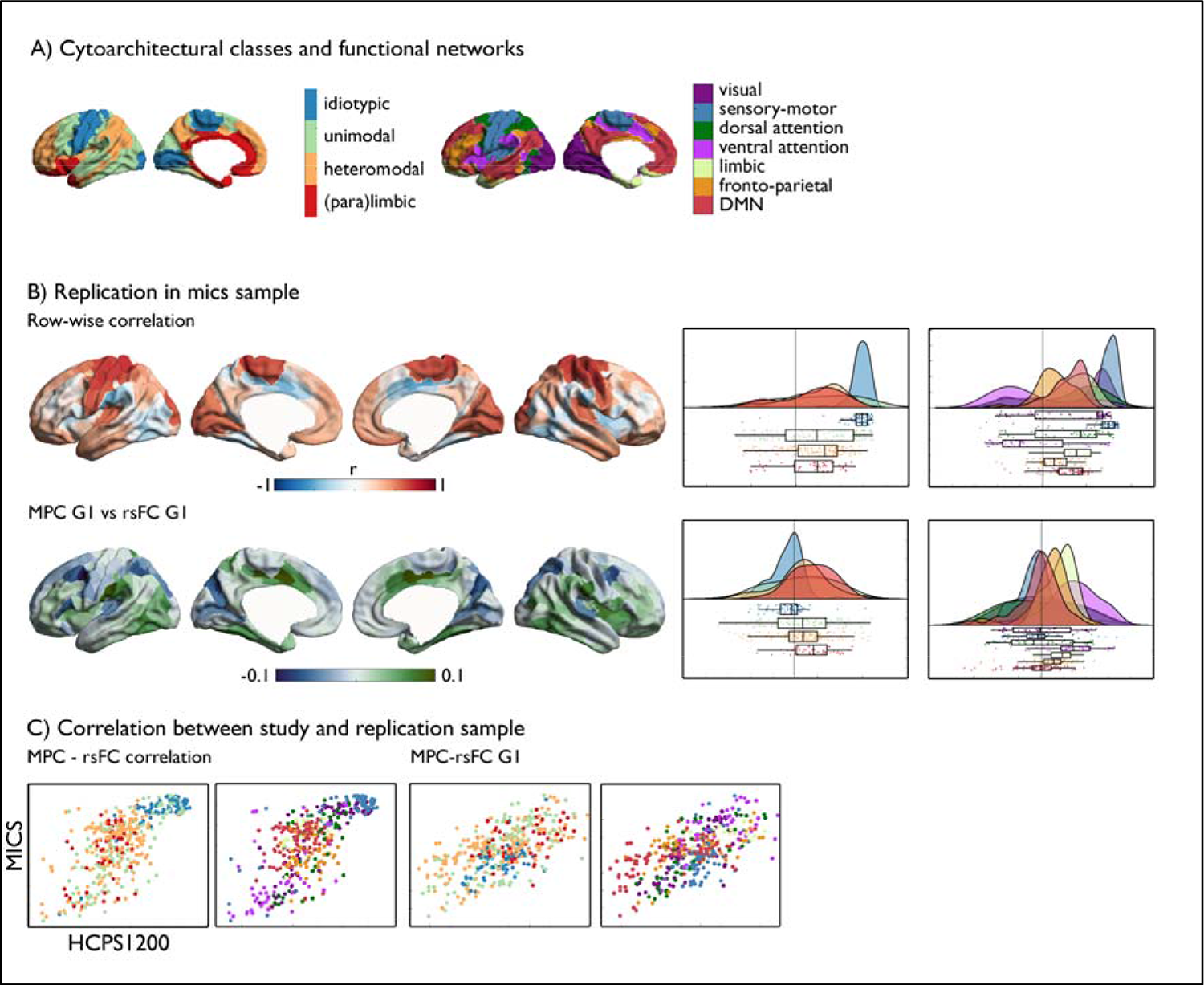
Replication sample MICS. **A**) Cytoarchitectural class ^49^, and functional networks ^27^; **B**) Replication in the mics sample; *upper:* row-wise correlation and associated raincloud plots in cytoarchitectural class (left) and functional networks (right); *lower:* difference between gradients and associated raincloud plots in cytoarchitectural class (left) and functional networks (right); **C**) Correlation between HCP 1200 and mics sample of MPC-rsFC edge-level correlation (left) and MPC-rsFC principal gradient difference (right), colored by cytoarchitectural class and functional network.

**Supplementary Figure 2.**
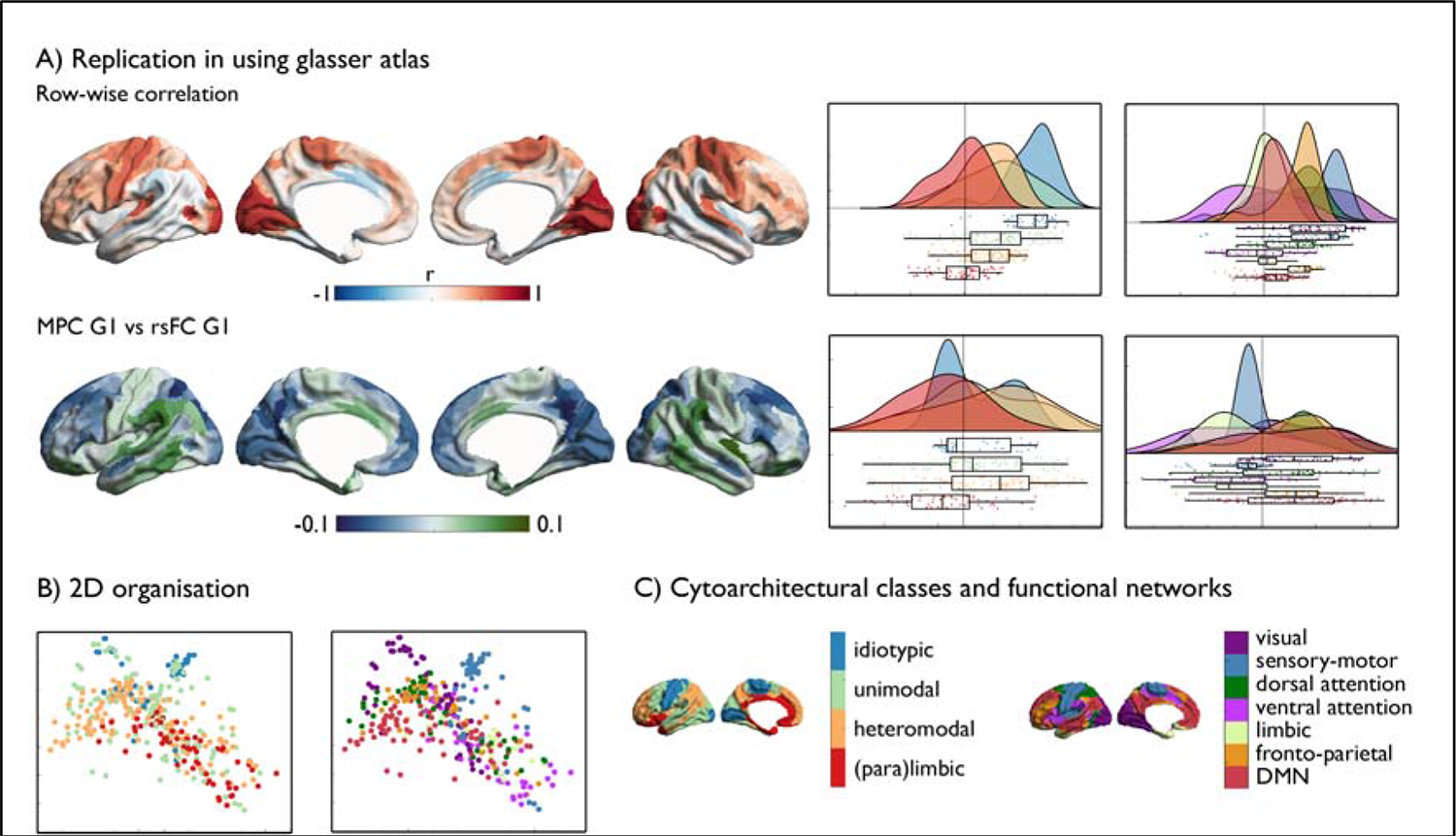
Replication in HCP using the Glasser atlas. **A**) Replication using the Glasser parcellation; *upper:* row-wise correlation and associated raincloud plots in cytoarchitectural class (left) and functional networks (right); *lower:* difference between gradients and associated raincloud plots in cytoarchitectural class (left) and functional networks (right); **B**) 2D organization x-axis gradient difference and y-axis structure-function row-wise correlation, colored by cytoarchitectural class (left) and functional network (right); **C**) Maps of cytoarchitectural class ^49^, and functional networks ^27^.

**Supplementary Figure 3.**
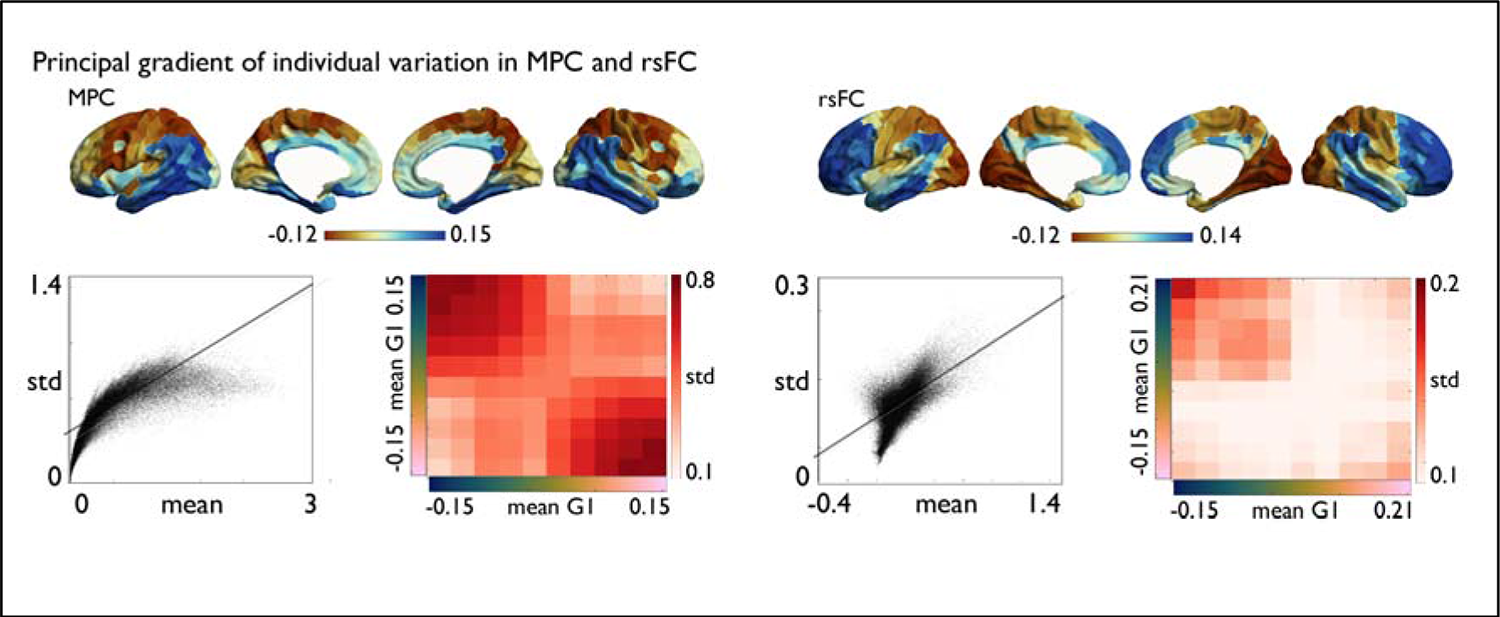
Principal gradient of individual variability. Principal gradient of individual variation (std) in MPC and rsFC, *lower panel left:* node-wise correlation between mean and std, and std along the mean gradient; *lower panel right:* node-wise correlation between mean and std, and std along the mean gradient

**Supplementary Figure 4.**
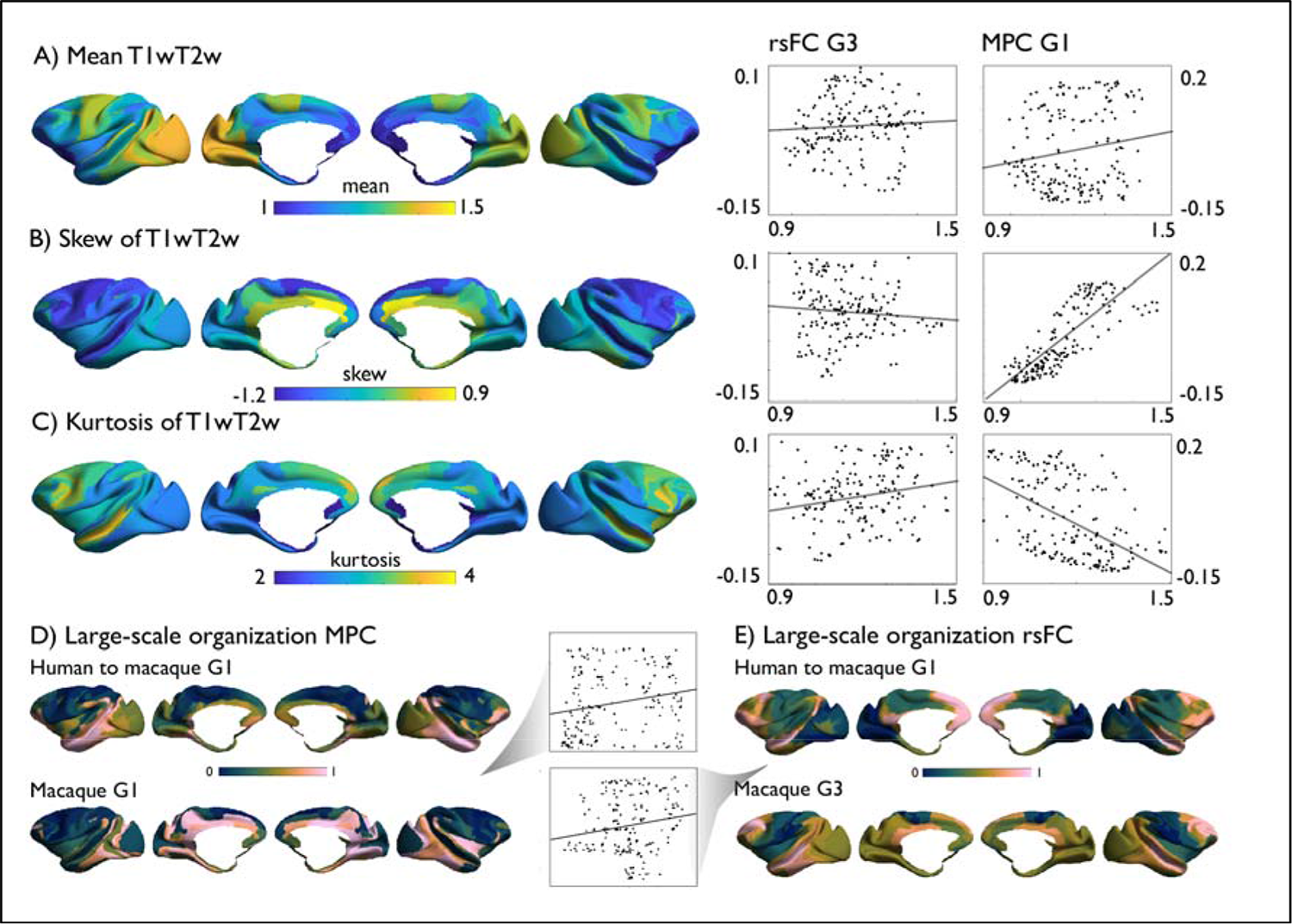
Macaque gradient validation. Validation of macaque MPC and rsFC gradients, using **A-C**) mean T1wT2w, skew, and kurtosis; **D**) Large-scale organization of MPC in human-to-macaque (upper) and macaque (lower) and its correlation; **E**) Large-scale organization of rsFC in human-to-macaque (upper) and macaque (lower) and its correlation.

## SUPPLEMENTARY TABLES

**Supplementary Table 1.**
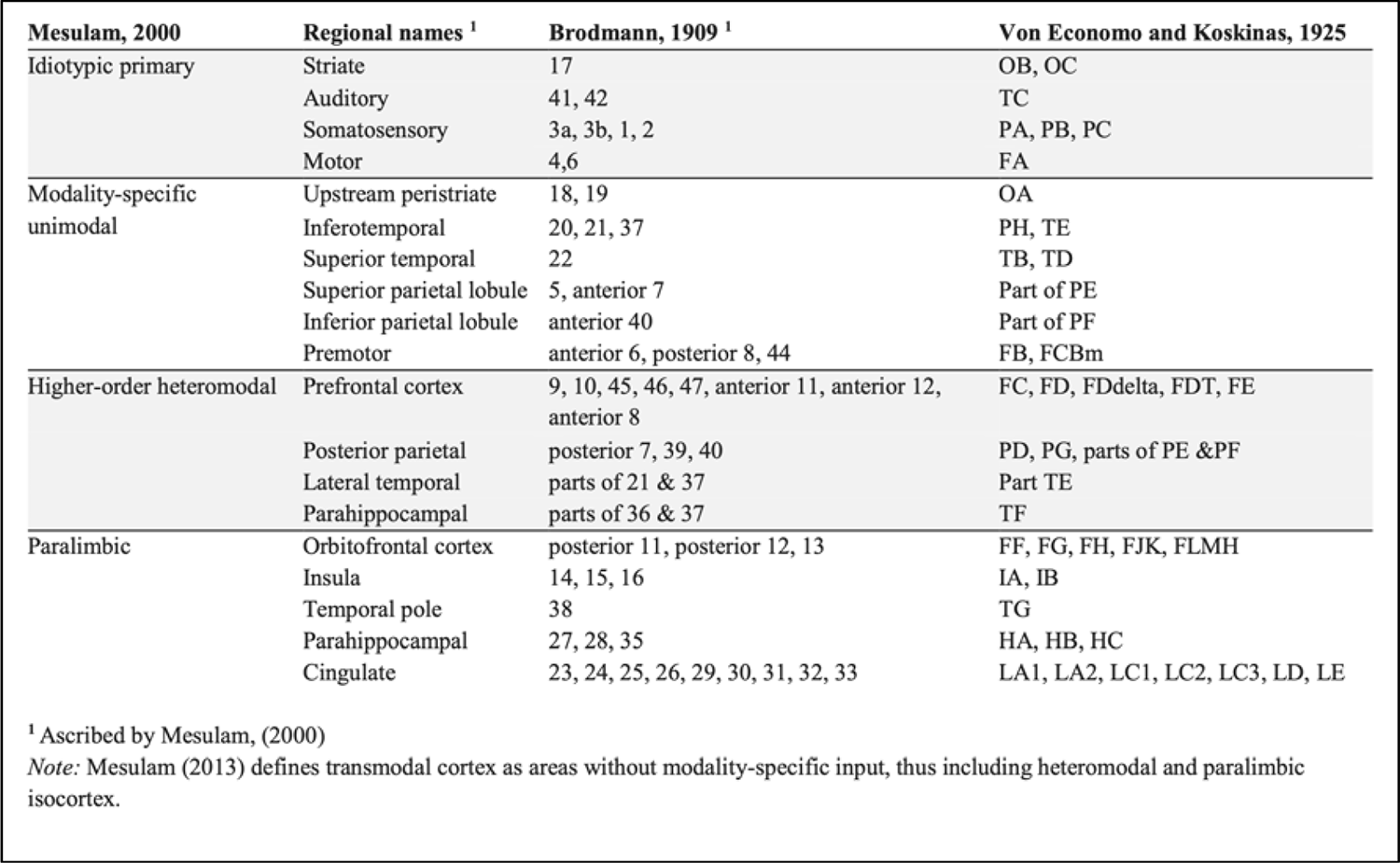
Replication of table in Paquola, 2019 of Mesulam classes nomenclature.

**Supplementary Table 2.**
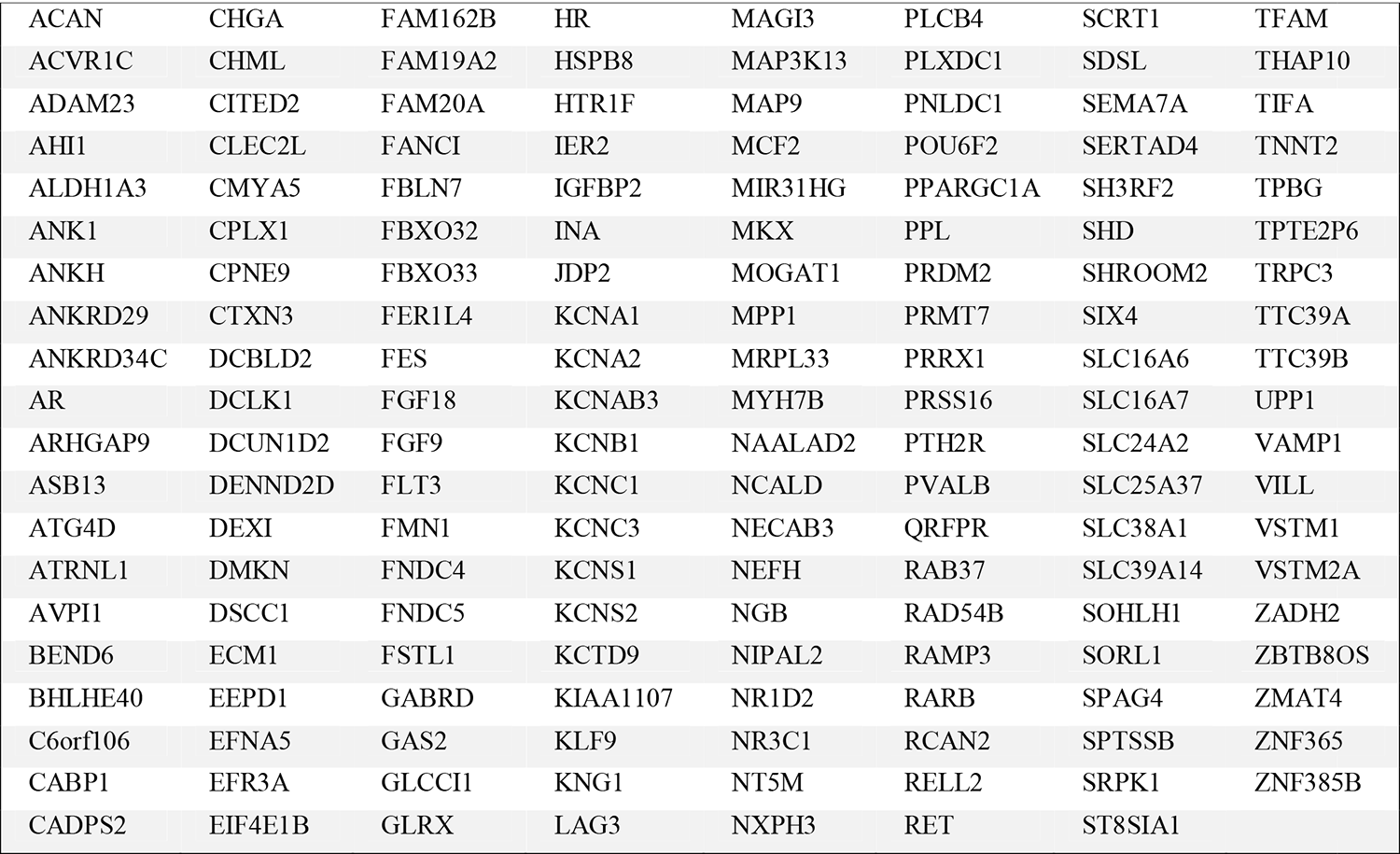
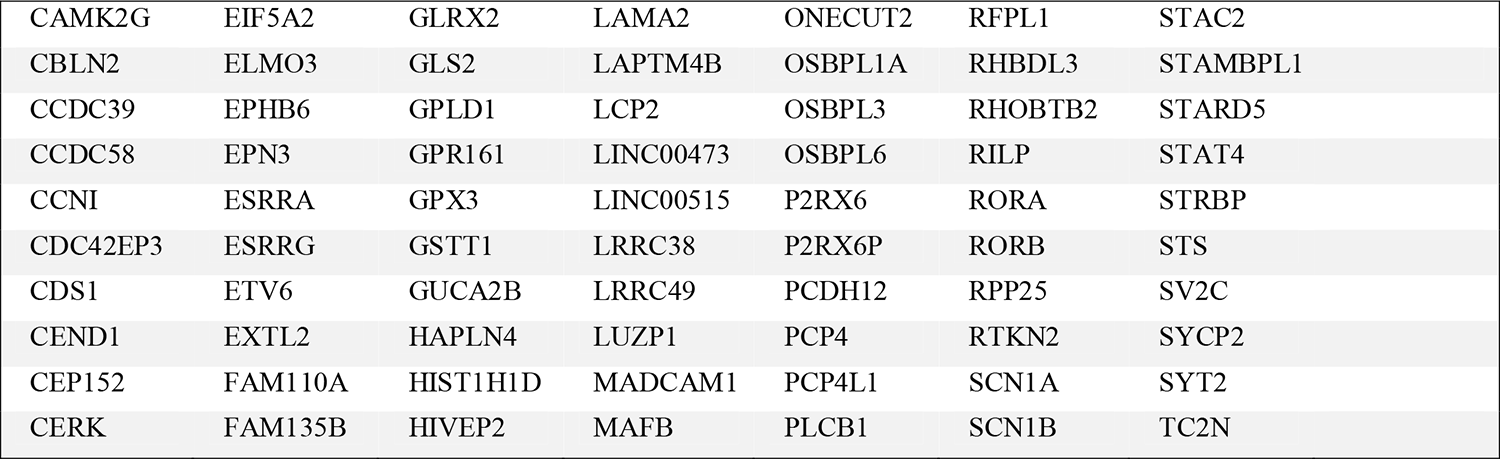
**rsFC>MPC.** List of genes that showed an increase along the gradient from rsFC to MPC in humans, r>0.5

**Supplementary Table 3.** **MPC>rsFC.** List of genes that showed an increase along the gradient from MPC to rsFC in humans, r>0.5.

|  |  |  |  |  |
| --- | --- | --- | --- | --- |
| ANKRD50 | FKBP1A | LINC00260 | PLXNC1 | SSTR1 |
| ANKRD6 | FNBP1L | LOC100129291 | PNMT | SULF2 |
| ASCL2 | GABRA5 | LOC642852 | PON3 | SYT17 |
| ATOH7 | GABRB1 | LOXL1 | PPM1M | TEKT2 |
| ATP2B4 | GABRE | LRRC36 | PPP4R4 | THRA |
| B9D1 | GLOD4 | LRRC3B | PRKCD | TLL1 |
| BIRC3 | GMFB | LUZP2 | PRKCG | TMEFF2 |
| CADM1 | GNG10 | LXN | PSD3 | TMEM159 |
| CAMK2D | GNG2 | MACROD2 | PTCHD1 | TMEM200A |
| CD24 | GNG4 | MAPK1 | PTGER3 | TNFRSF11A |
| CFD | GPC4 | MARCKS | PTPRA | TUBB2A |
| CLEC4G | GPD2 | MESP1 | PTPRR | UCHL3 |
| COCH | GPR160 | MMD | PYDC1 | WDR66 |
| CPE | GRIK2 | NANOS1 | RAB27B | WDR86 |
| CPNE6 | GRP | NCAM2 | RAPGEF4 | XKR4 |
| CPNE7 | HPCAL4 | NECAB2 | RAVER2 | YPEL1 |
| CTXN1 | HSPB2 | NKAIN4 | RILPL2 | ZCCHC12 |
| CXorf57 | HTR1A | NOV | RNF150 | ZCCHC17 |
| DACH2 | HTR2C | NPPA | RRP7A | ZCCHC18 |
| DHDH | IL13RA2 | NR2F2 | RSPH9 | ZNF436 |
| DIAPH2 | IQCJ | NTSR1 | SCGN |  |
| DIRAS3 | ISG15 | NUDT11 | SCN3B |  |
| DOK6 | ISOC1 | OPRM1 | SCN9A |  |
| DPYSL3 | KCNA4 | PCSK5 | SHF |  |
| DYDC2 | KCNG1 | PDYN | SLA |  |
| EFCAB1 | KCNN3 | PEA15 | SLC16A2 |  |
| F12 | KCTD12 | PGAP1 | SLC17A8 |  |
| FABP7 | KCTD4 | PGRMC1 | SLIT1 |  |
| FAM171B | KLHL13 | PID1 | SLIT3 |  |
| FAM71F1 | KLK10 | PKIA | SMARCD3 |  |

**Supplementary Table 4.**
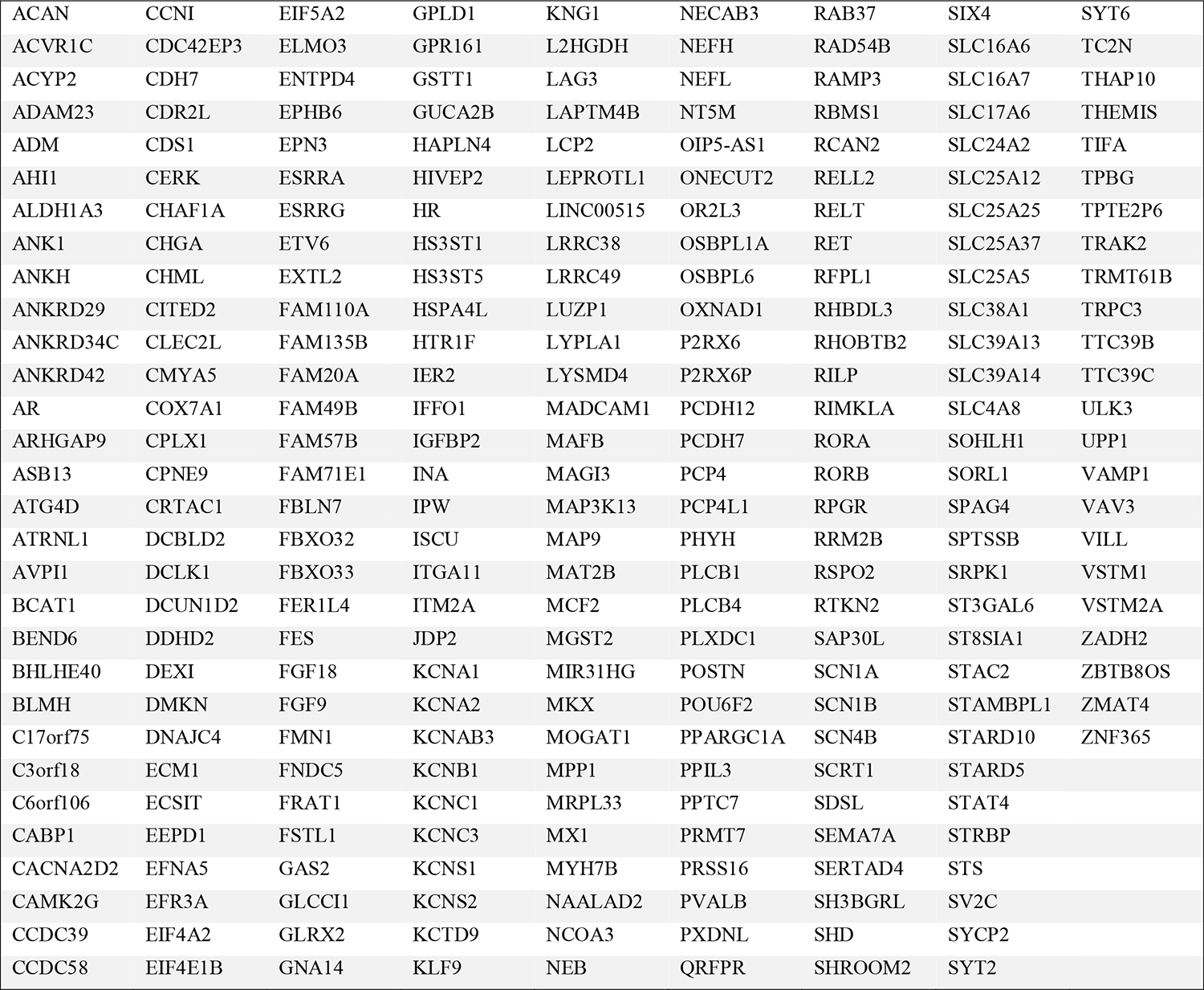
**Coupling** List of genes that showed an increase with increased coupling in humans, r>0.5.

**Table 5.**
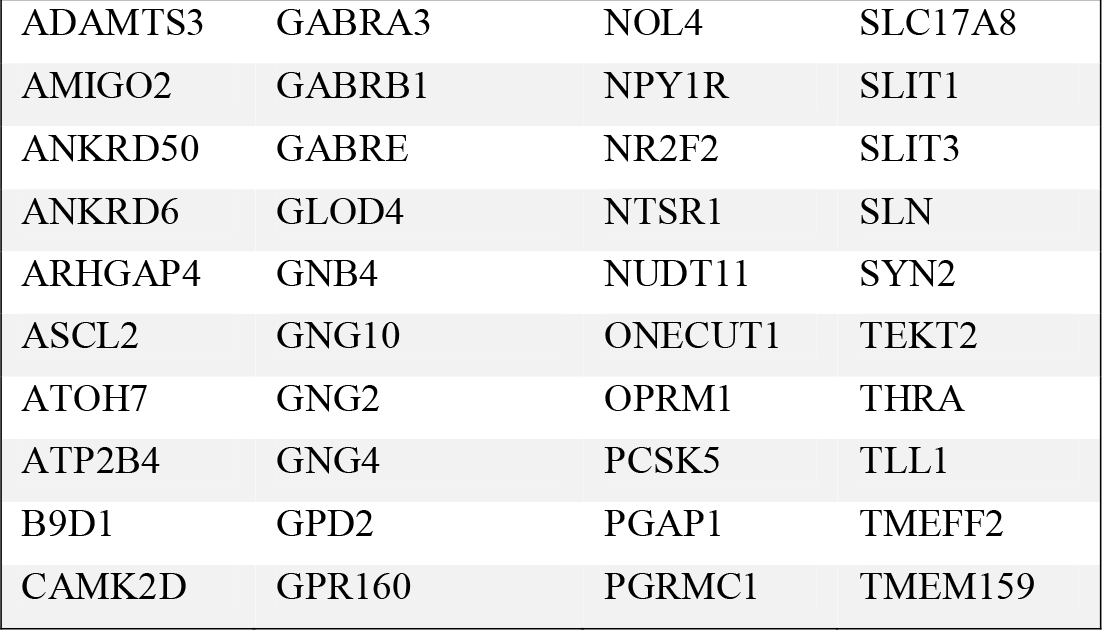
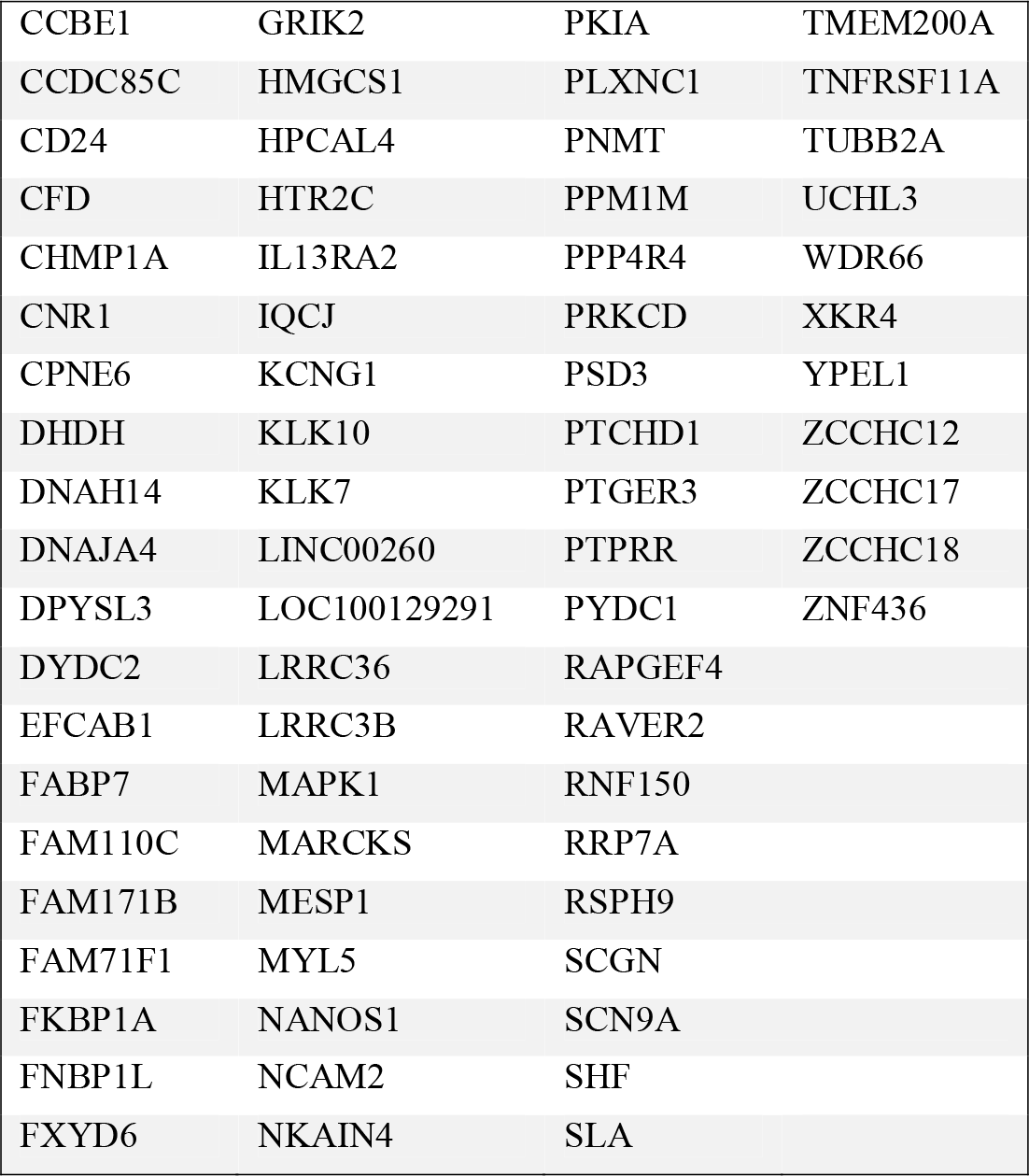
Uncoupling. List of genes that showed a decrease with increased uncoupling in humans, r>0.5.

